# A computational method for detection of structural variants using Deviant Reads and read pair Orientation: DevRO

**DOI:** 10.1101/094474

**Authors:** Shumaila Sayyab, Nima Rafati, Miguel Carneiro, Hervé Garreau, Göran Andersson, Leif Andersson, Carl Johan Rubin

## Abstract

**Background:** Next generation sequencing (NGS) technology has made it possible to perform high-resolution screens for structural variants. Computational methods for detection of structural variants utilize paired-end mapping information, depth of coverage, split reads, or some combination of such data. The available methods are particularly designed to detect structural variants in single genomes or multiple genomes in a pairwise manner. The aim of this study was to develop a bioinformatics pipeline for detection of large structural variants using multiple pooled populations.

**Results:** Here we describe the method “DevRO”, developed to enable identification of structural variants using short insert paired-ends and long-range mate-pairs. DevRO uses paired-end mapping information from both types of libraries for identification of inversions, deletions and duplications followed by read depth information to screen for copy number variants. DevRO can detect structural variants in multiple populations without the need for pairwise comparisons. It uses a combined approach based on (*i*) paired-end mapping and (*ii*) depth of coverage that gives power to the study as compared to traditional methods that are based on either of these. DevRO is also designed to detect deletions in the reference assembly, which is an added functionality as compared to available methods.

**Conclusion:** We report a bioinformatics pipeline “DevRO” for detection of structural variants using paired-end mapping and depth of coverage methods tested on sequencing reads from multiple pooled rabbit populations. This method is useful when large numbers of populations have been re-sequenced as compared to traditional methods that can detect structural variants in a pairwise manner.

## Background

In human genetics a focus has been to identify rare structural variants associated with disease whereas in animal genetics as well as in evolutionary biology it is of considerable interest to identify loci under positive selection. Large structural variants (SV) are an important form of genetic variants that frequently underlie phenotypic variation (Andersson, 2013; Weischenfeldt *et al.,* 2013). SV refers to copy or dosage changing variants called copy number variations (CNVs which include deletions, insertions and duplications (Redon *et al.,* 2006)) or dosage neutral variants like inversions and translocations. In humans, approximately 1.2% of a single genome differs from the reference genome as regards CNV genotypes (Pang *et al.,* 2010). The average size of SVs in the human genome is ~8kb and they range from 50bp to large structural events (Alkan *et al.,* 2011). In the recent past, several studies indicated that SVs have been associated with a variety of human diseases (Yang *et al.,* 2013; McCarroll & Altshuler, 2007; Stranger *et al.,* 2007). Whereas in livestock genomes, research in genome-wide CNV identification of various domestic animals showed their importance either in disease association (Olsson *et al.,* 2011), phenotypic changes (Jia *et al.,* 2013; Imsland *et al.,* 2012; Rubin *et al.,* 2010) affecting different traits, association with breed-specific differences in adaptation, health, and production traits (Bickhart *et al.,* 2012) and adaptation to starch rich diet in dogs after domestication (Axelsson *et al.,* 2013).

Before the development of next generation sequencing, structural variants were discovered by either hybridization methods (aCGH) in which the relative probe hybridization intensities differ between two compared genomes (Pinkel *et al.,* 1998) or using Hapmap data and SNP arrays measuring the intensities of probe signals at SNP loci (International HapMap *et al.,* 2010). However, the power was limited because of the size and breakpoint resolution of the predicted SV due to the density of SNP arrays. Sanger sequencing of paired reads was used as an alternative to the above-mentioned methods to detect CNVs, inversions and translocations with high accuracy and resolution at the expense of time and cost. Today next generation sequencing technologies (NGS) can generate a large amount of sequence data for instance by whole genome sequencing at a fraction of the cost and time. Several methods have been developed to enable SV detection from NGS data but each method has some limitations. In general, there are four categories of methods to detect SVs using NGS data 1) Depth of Coverage (DoC); 2) Paired-end mapping (PEM); 3) Split read (SR) and 4) Assembly (AS) based methods (Alkan *et al.,* 2011).

The assumption of depth of coverage based methods (*e.g.* CNVseq and CNVnator) is that the coverage is uniform *i.e.* the number of reads mapped to a region are assumed to follow a Poisson distribution, however these methods are unable to detect inversions and translocations (Abyzov *et al.,* 2011; Xie & Tammi, 2009). Paired-end mapping methods (*e.g.* DELLY and Breakdancer) use the information of paired reads and their orientation. The insert size is used to detect insertions and deletions, although the size of CNVs detected is limited by insert size of the libraries used (Rausch *et al.,* 2012; Chen *et al.,* 2009). Split read methods (*e.g.* Pindel) use the information of anchored reads to identify the breakpoint locations while assembly based methods (*e.g.* SOAPdenovo) are based on a denovo assembly (Li *et al.,* 2010; Ye *et al.,* 2009). Today, several tools have been developed that use a combination of the data available (*e.g* GenomeSTRiP, SVDetect) (Handsaker *et al.,* 2011; Zeitouni *et al.,* 2010).

Most of the available methods either use data from a single sample (CNVnator) or make pairwise comparisons between samples (CNVseq). The complexity arises when there are multiple groups (*e.g.* group1 comprising n populations and group2 comprising m populations) then detection of SVs needs further pairwise comparisons at the downstream level, making it difficult to attribute SVs to a specific group. Another complicating problem arises if the reference genome carries a deletion of a segment or if the resequenced individual/population carries an insertion not present in the reference, because the reads from this region cannot be mapped to the reference genome. This may for instance occur when comparing domestic animals with their wild ancestors, if a DNA segment has been deleted during the domestication process. Clearly, genome sequences from more individuals are needed to define whether a certain SV allele is derived or ancestral.

The aim of this study was to develop a tool based on deviant read and read orientation (DevRO) information using both paired-end mapping and depth of coverage analysis that can detect (*i*) structural variants in multiple populations and (*ii*) detect deletions in a reference assembly compared with individuals carrying a non-deleted allele.

## Results and Discussion

### Detection of structural variants using whole genome resequencing data

Here we used BAM files (Li *et al.,* 2009) as input to DevRO VariantCaller. The VariantCaller step generated raw variants calls from multiple individuals. In DevRO VariantCaller, we scan the entire genome for PEM signatures (Figure 1) in windows of 1 kb. For each locus we store the information of counts of discordant reads in each population, median mate position for the discordant reads within 2 standard deviations of the mapping distance between mate pair reads, forward and reverse median position of the anchored (singleton) reads and their counts in each population, soft clipped read counts in each population along with forward and reverse strand clipped positions and concordant read counts (Figure 2, Figure S1). This information is further processed at the VariantParser step to calculate the fraction of deviant reads in each group. This information is used to identify loci showing significant differences between groups. The CNVs detected using VariantParser are annotated and scored using the DoC information from paired-end sequencing data.

**Figure 1.**
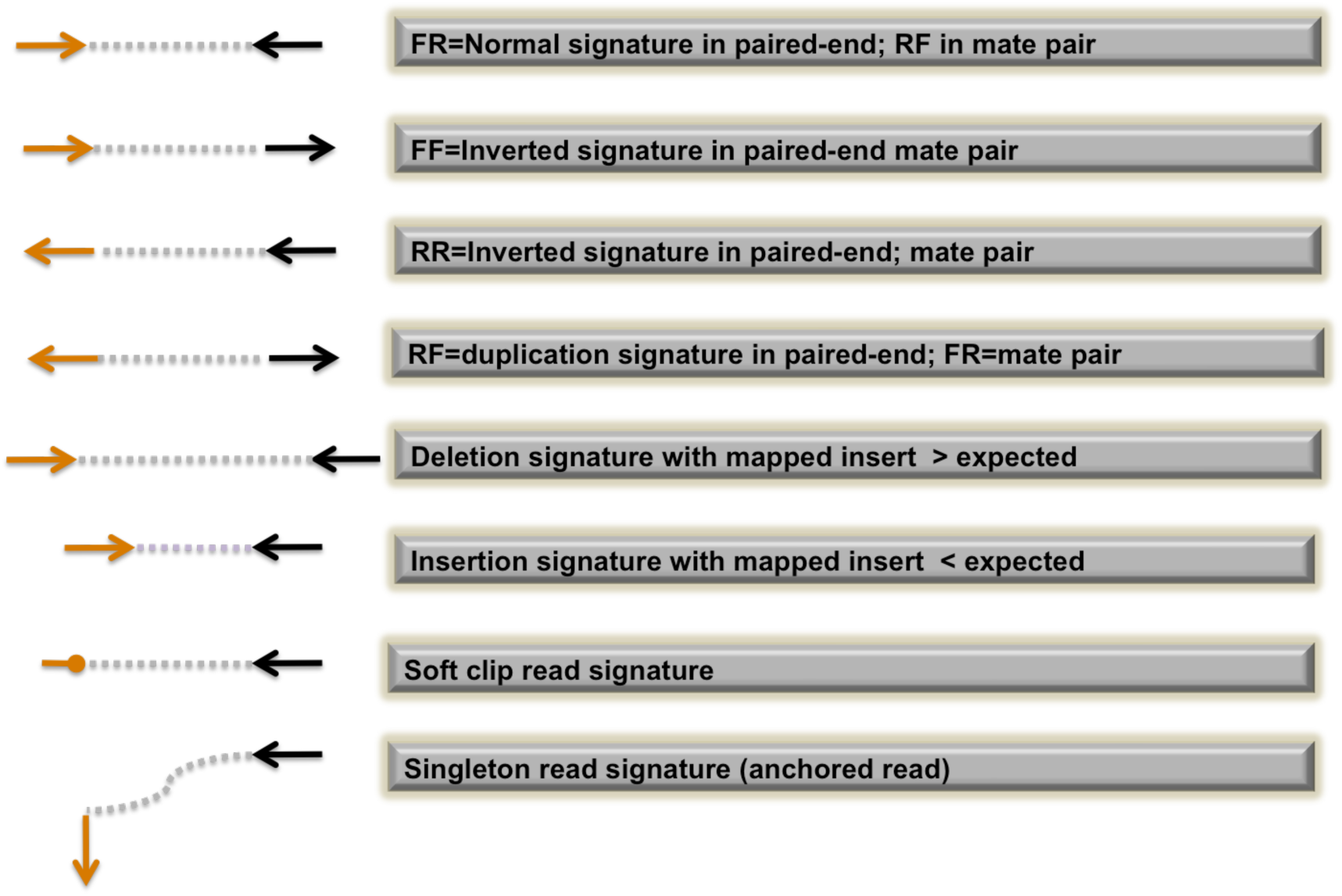
Description of signatures used in the analysis of paired end mapping data. Forward read marked as F and Reverse read marked as R.

**Figure 2.**
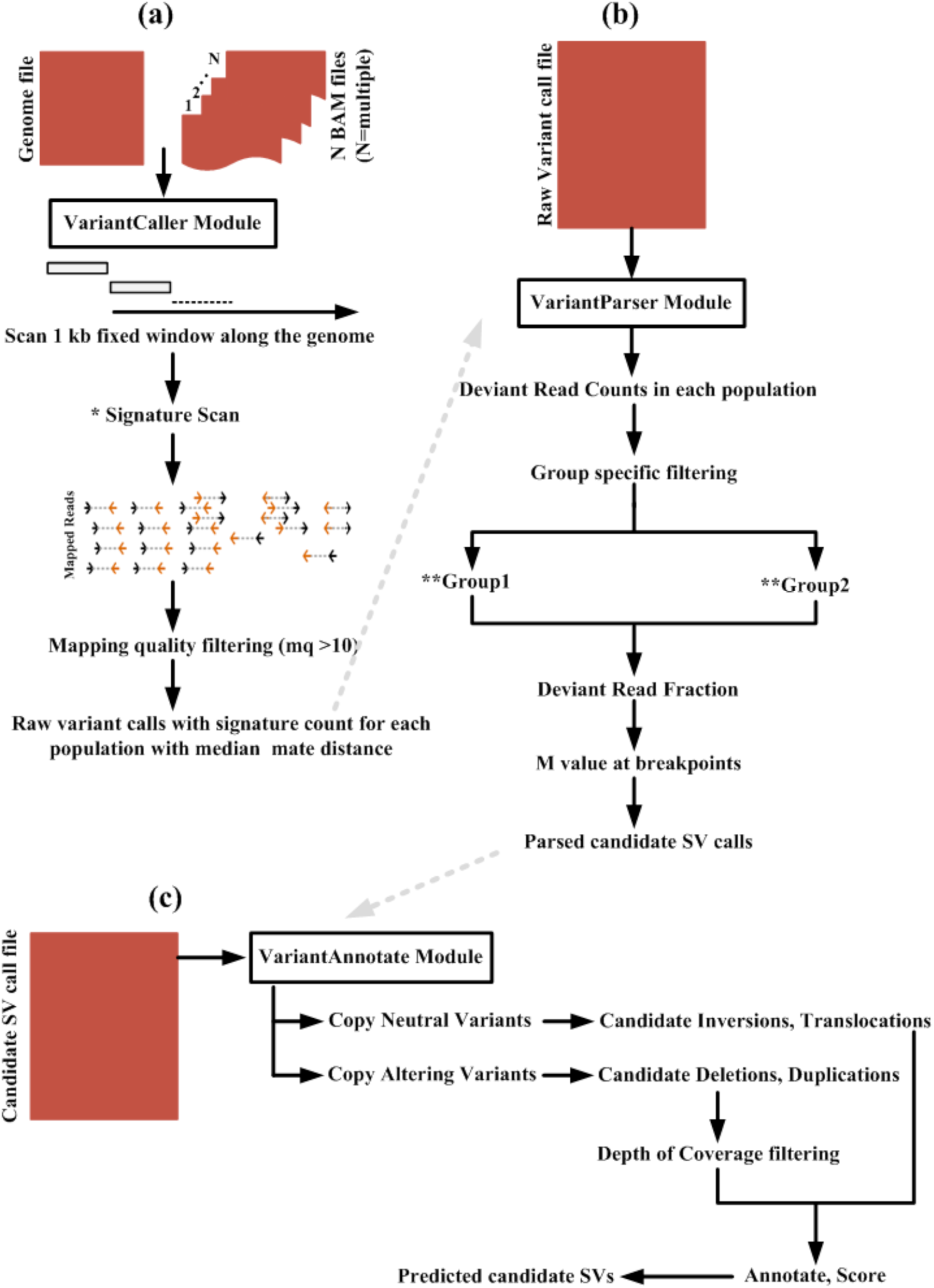
Overview of DevRO SV screen. **a)** Steps in VariantCaller module; *For detailed workflow see Figure S1 **b)** Steps in VariantParser; ** Group1 and Group2 represents two groups in test data and **c)** VariantAnnotate.

As a test case we used two data sets. 1) Mate pair (MP) data generated from two wild and two domestic rabbits (average insert size of 4.5 kb and average coverage of 3x) for PEM signatures (Figure 1). 2) Rabbit paired-end (PE) sequencing data from pooled samples of wild and domestic rabbit populations with an average coverage of 10x per population (Carneiro *et al.,* 2014) was used for DoC information. For this particular test case, DevRO uses information of two groups (wild and domestic rabbits) to find SVs with significant frequency differences between the two groups. However, it is not limited to this scenario and can take any two groups (*e.g.* case and control).

This resulted in identification of candidate deletions, duplications (Table S1) and preliminary results of inversions present at a relatively high frequency in either group 1 or group 2. Figure 3 and Figure S2 shows few examples of candidate duplication and deletions predicted by DevRO. Further improvements utilizing for example base quality information of deviant reads, and mapping of breakpoint read sequences on the reference genome will be added to later versions of DevRO.

**Figure 3.**
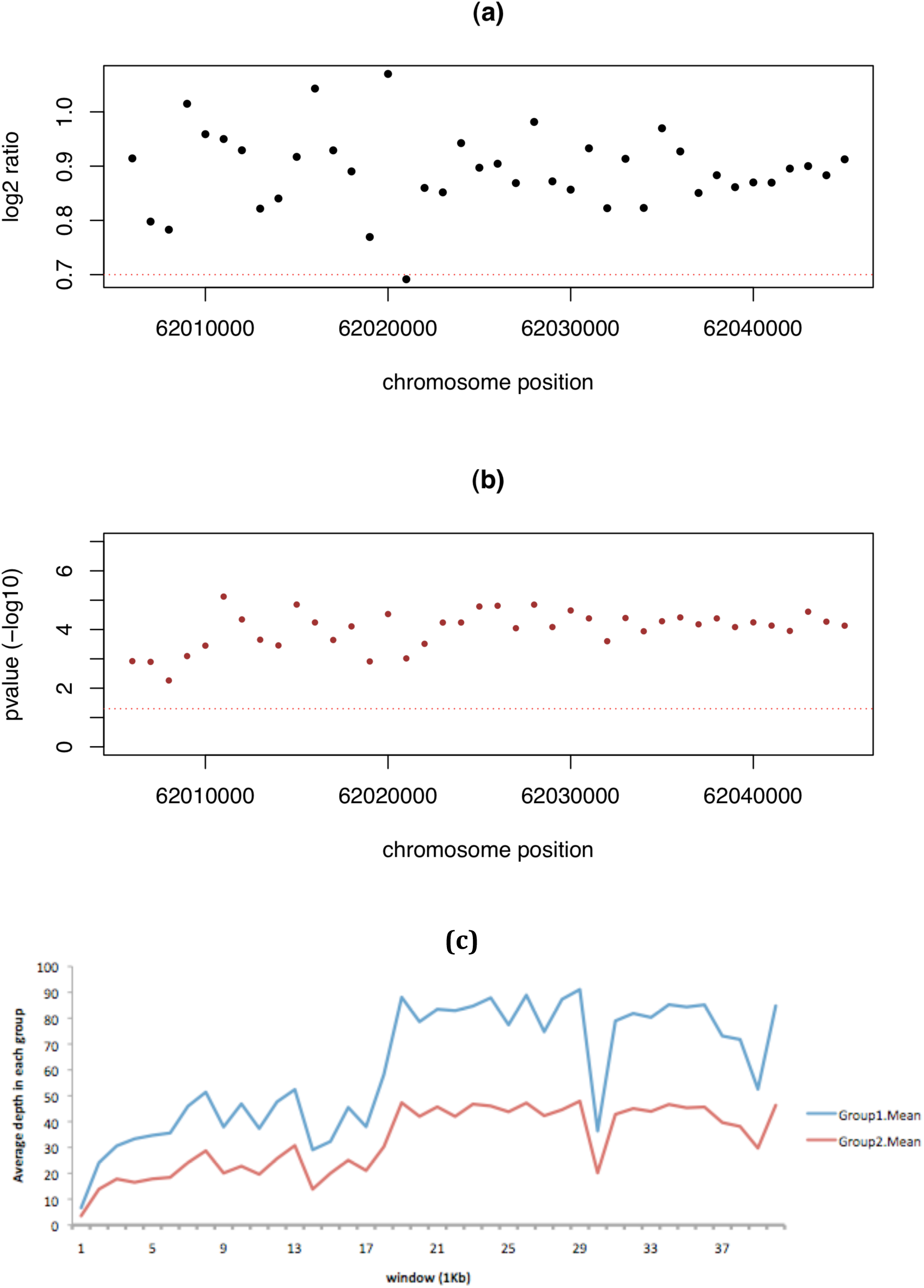
Example of Candidate duplication detected by DevRO supported by 16 discordant reads from MP data. **(a)** Mvalue plotted (black dots), with dotted horizontal red line threshold 0.7 **(b)** pvalue plotted with (brown dots) showing the significant difference in depth between group 1 and group2. **(c)** Mean depth plotted for each group in windows of 1kb size using PE data.

### Condordance with SVDetect and Breakdancer

SVDetect (Zeitouni *et al.,* 2010) is also based on the combined use of PEM and DoC data for detecting SVs in multiple samples. Breakdancer (Chen *et al.,* 2009) is based on the PEM approach for detection of SVs (deletions, inversions and translocations). The main difference between DevRO and these tools is that DevRO, as opposed to SVDetect and Breakdancer, does not require pairwise comparisons of groups (Table 1). We ran SVDetect and Breakdancer using the rabbit mate pair data in order to assess concordance with DevRO results (Table 2). The number of inversions detected was 178 and 281 for SVDetect and DevRO respectively, with 248 overlapping inversions using bedtools feature “reciprocal fraction overlap” of 0.7. Table S2 shows the preliminary list of candidate inversions detected by DevRO not in SVDetect list. One possible reason why SVDetect detected fewer inversions could be it is based on pairwise comparisons as opposed to DevROs group level contrasts. The overlap between SVDetect and DevRO was 500 and 391 for deletions and duplications, respectively (Table 2). However, the overlap between Breakdancer and DevRO showed big differences 13 and 43 for inversions and deletions, respectively. One possible reason for this big difference in deletions could be due to the combined use of PEM and DoC in DevRO while only using PEM in Breakdancer for detection of SVs.

**Table 1.** Summary comparison of DevRO with available SV softwares.

|  |  | **SVDetect** | **Breakdancer** | **DevRO** |
| --- | --- | --- | --- | --- |
| Input | *BAM | √ | √ | √ |
| Output | INV | √ | √ | √ |
|  | INS | √ | √ | √ |
|  | DEL | √ | √ | √ |
|  | DUP | √ | – | √ |
|  | TRANSL | √ | √ | √ |
|  | Del-Ref | – | – | √ |
| Group-wise Comparisons | Parser | – | – | √ |
| Visualization | Output plots | √ | – | √ |
| Methods |  | PEM+DoC | PEM | PEM+DoC |

**Table 2.** Summary of strucrural variants detected by SVDetect, DevRO and Breakdancer tools.

| Tool <sup>a</sup> | Inversions | Deletions | Duplications <sup>b</sup> |
| --- | --- | --- | --- |
| SVDetect (SVD) | 178 | 702 | 408 |
| DevRO | 281 | 796 | 462 |
| Breakdancer (BD) | 289 | 64 | NA |
| OL_SVD-DevRO | 248 | 500 | 391 |
| OL_BD-DevRO | 13 | 43 | NA |
| OL_SVD-BD | 65 | 62 | NA |

<sup>a</sup> Overlap as "OL", Breakdancer tool as "BD" and SVDetect tool as "SVD"
<sup>b</sup> Breakdancer only predicts inversions and deletions and unknown, Here Unknown not taken into account. NA=not available.

### Deletions in the rabbit reference genome

A unique feature with DevRO is that none of the previously described tools are designed to search for deletions present in the reference assembly. This is of particular interest in draft assemblies as well as in domestic animals when the reference genome is based on a domesticated individual, since this makes it challenging to identify regions of the genome that may have been lost during domestication. In rabbits, is based on a domesticated individual, as is the case for all other domesticated species except the chicken. One of the aims with the development of DevRO was to identify deletions present in a reference assembly; such deletions may be due to assembly errors or because the individual used for the assembly carry one or more deletions. In our case, the pileups of singleton reads (anchored reads whose pairs are unmapped) using data from wild rabbits were used to identify candidate loci. These were further narrowed down using the median positions for forward and reverse singletons and soft clipped positions. This resulted in a preliminary candidate list of breakpoints of deletions in the reference genome. These putative breakpoints of deletions were further used to extract the hanging reads. By combining the anchored and hanging read sequences (which represents one mapped read and one unmapped read), BLAT mappings were conducted to the human reference genome in order to investigate whether the unmapped reads corresponded to a human sequence homologous to the rabbit locus as this would indicate that the reference carried a deletion of evolutionary conserved sequences. Further improvements are needed to conduct *de novo* assembly of the singletons and unmapped reads at breakpoints. Further work is also required to experimentally validate these putative deletions.

In order to test DevRO for detection of deletions in reference assembly, we simulated deletions in the rabbit reference assembly (Table S3). Here we deleted ten regions ranging in size from 1 kb to 70 kb on chromosome 1 with repeats (repeat information extracted from UCSC Santa Cruze Browser) flanking the breakpoints or overlapping them. DevRO successfully identified nine of the simulated deletions but was unable to detect one of the regions (Tables S3). This region showed a mappability of 0.008 at the 5’ end of the breakpoint indicating a highly repetitive region.

## Conclusions

Here we report a bioinformatics pipeline “DevRO” for detection of structural variants using paired-end mapping and depth of coverage analysis and tested it on rabbit data. It has an added routine for detecting deletions present in a reference genome. This method will be useful when large numbers of populations are re-sequenced as compared to other frequenctly used methods it is designed to detect structural variants in pairwise comparisons of groups.

## Methods

The input of DevRO is a set of aligned MP or PE reads in SAM/BAM format (Li *et al.,* 2009). All input BAM files from multiple populations are analyzed jointly, to avoid pairwise comparisons in the end.

The pipeline DevRO consists of three modules 1) VariantCaller 2) VariantParser and 3) VariantAnnotate that are used to detect inversions, deletions and duplications when comparing two or more populations as well as deletions in a reference genome seq.

### Variant Caller

We used BAM files (Li *et al.,* 2009) from MP data as an input to DevRO VariantCaller. In this step, we scan the genome in windows of 1kb to search for PEM signatures using discordant reads (Figure 1). The discordant or deviant reads have (***I***) abnormal mapping distance in comparison to average mapping distance, the threshold for declaring a PEM as deviant is that it deviates more than three standard deviations from the mean which is a signature for deletion or insertion, (***II***) abnormal relative mapping orientation (in case of duplication or inversion) as shown in Figure 1. The information of the mate orientation strand for the left or right clipped reads to identify breakpoints. Together with counts and position of discordant reads we also recorded the counts of anchored or singleton reads and soft clip reads using bitwise flags and CIGAR (Li *et al.,* 2009) given in alignment files (BAM) as the method described in SAMtools (Li *et al.,* 2009). For duplications, we store all those loci where at least one duplication read signature is observed in any population, and record the information of deviant/discordant and normal read counts for such loci in each population, median positions are calculated using the mates (discordant reads). This step gives us the raw calls for loci with read counts in each SV category for all the populations in question. The following filters were used during this step: mapping quality for discordant reads >=10; same criteria is applied for each SV type for which the VariantCaller is run. Each analysis is done separately for duplications, inversions and deletions. This means that when we are running the script for the duplication scan and we come across inverted loci having no duplication reads then these loci will not be reported in output of duplications call but will be reported in inversion calls. In contrast, if we find inverted reads where we have duplicated reads also, the loci will be stored with the information of read counts for duplications and inversions. The same principle applies for the scan of inversions.

### Variant parser

The purpose of this module is to extract only those loci where there is differentiation between the two groups analyzed (domestic and wild rabbits in the data analyzed in this study). The input is the result of Variant Caller raw calls obtained in the above step. In this step we calculate the fraction of abnormal reads in the two groups and only extract the loci where we see a significant difference between the two groups.

The formula for calculating the fraction of reads consistent with duplication in group 1 is as follows:

> frac_dup = A1_dup/(A1_dup+A2_dup), where A1 and A2 represents two groups.

The same type of formula is used to calculate the fraction for all discordant or deviant reads.

### Variant Annotate

The predicted CNVs from VariantParser are given as an input to Variant Annotate. We calculate chisq test for each candidate using normalized deviant read counts in each group (A1 and A2) and the total read counts. The purpose of this step is to score and annotate the loci identified in the variant parser step by using pvalue and information of depth of coverage for CNVs using paired-end data (method explained in Carneiro et al., 2014). For each group the average depth is extracted in the predicted regions and an M-value (log2 fold change) is calculated. This step gives CNVs that show significant allele frequency difference between two groups (absolute M-value >=0.7).

For visualization, log2 fold change is plotted for each candidate CNV in R. The breakpoints for inversions are shown using the positions of mate pair. We next annotate the candidates using information for genes, repeats, gaps or custom annotations. Finally, candidate SVs are scored as alpha, beta and gamma by using the following criteria for all SVs:

Alpha: SV with absolute M-values greater than 0.7 from MP and PE data and that fulfill the fraction check, where fraction check is at least 10% of deviant reads or signatures observed.

Beta: Below 10% and above 1% of deviant reads support with or without M-values. Gamma: without M-value support, with less than 1% deviant reads.

### Deletion in the reference genome assembly

For detecting deletions in the reference genome or insertions in the resequenced genome, the genome was scanned as in the VariantCaller step by using pileups of singleton (anchored reads) in non-reference populations with mapping quality greater than 10. Median position is calculated for both forward and reverse singleton reads with counts for discordant reads in each window. If multiple discordant reads are observed, then it is less likely that the singletons are due to deletion in the reference assembly and we discard that window in parser step. We further narrow down the region by making use of median positions of singletons and only allow 0.05% overlap between forward and reverse singletons (if any). Together with this information, soft clips (search for right and left clipped positions at breakpoints) are used to detect breakpoints of deletions in the reference assembly. In order to know what part may have been deleted in the reference genome, we also make use of BLAT to map the unmapped pairs of singletons at breakpoints to the human genome assembly to infer whether a DNA fragment observed in human (and in other species) is homologous to those unmapped reads. The size and expected gene content within the part missing in the reference assembly is inferred.

### Datasets

Four rabbits (two domestics and two wild from the *Oryctolagus cuniculus algirus* subspecies) were re-sequenced at 3x coverage using Illumina mate pair (2x50bp) with an average insert of 4.5kb. This dataset was used for paired-end mapping analysis. The depth of coverage analysis was done for the CNVs using the previously published paired-end sequencing data (Carneiro et al., 2014).

## Acknowledgements

The work was supported by the ERC grant BATESON to LA, by POPH-QREN funds from the European Social Fund and Portuguese MCTES [postdoc grant to M.C (SFRH/BPD/72343/2010)], by FEDER funds through the COMPETE program and by Portuguese national funds through the FCT – Fundação para a Ciência e a Tecnologia – (PTDC/CVT/122943/2010), by an EU FP7 REGPOT grant [CIBIO-New-Gen][286431], and by travel grants to M.C. (COST Action TD1101) and Higher Education Commission in Pakistan (support for S.S.). Computer resources were supplied by UPPNEX at Science for Life Laboratory.

## Supplementary Files

**Figuere S1.**
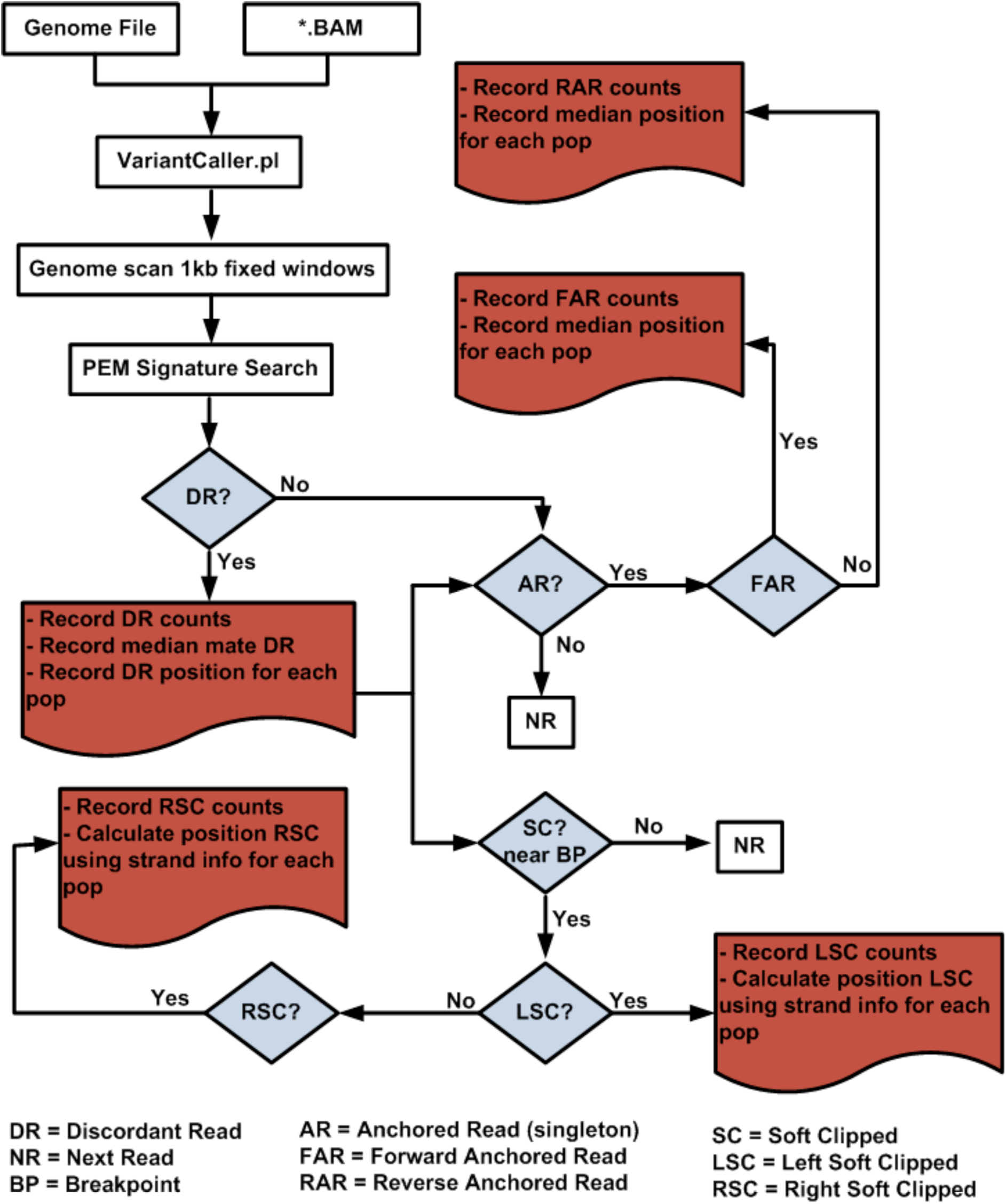
Detailed Flowchart of DevRO VariantCaller module. Discordant reads include inverted reads or duplicated reads or deletion or insertion signatures (see catalog of signatures, Figure 1).

**Figure S2.**
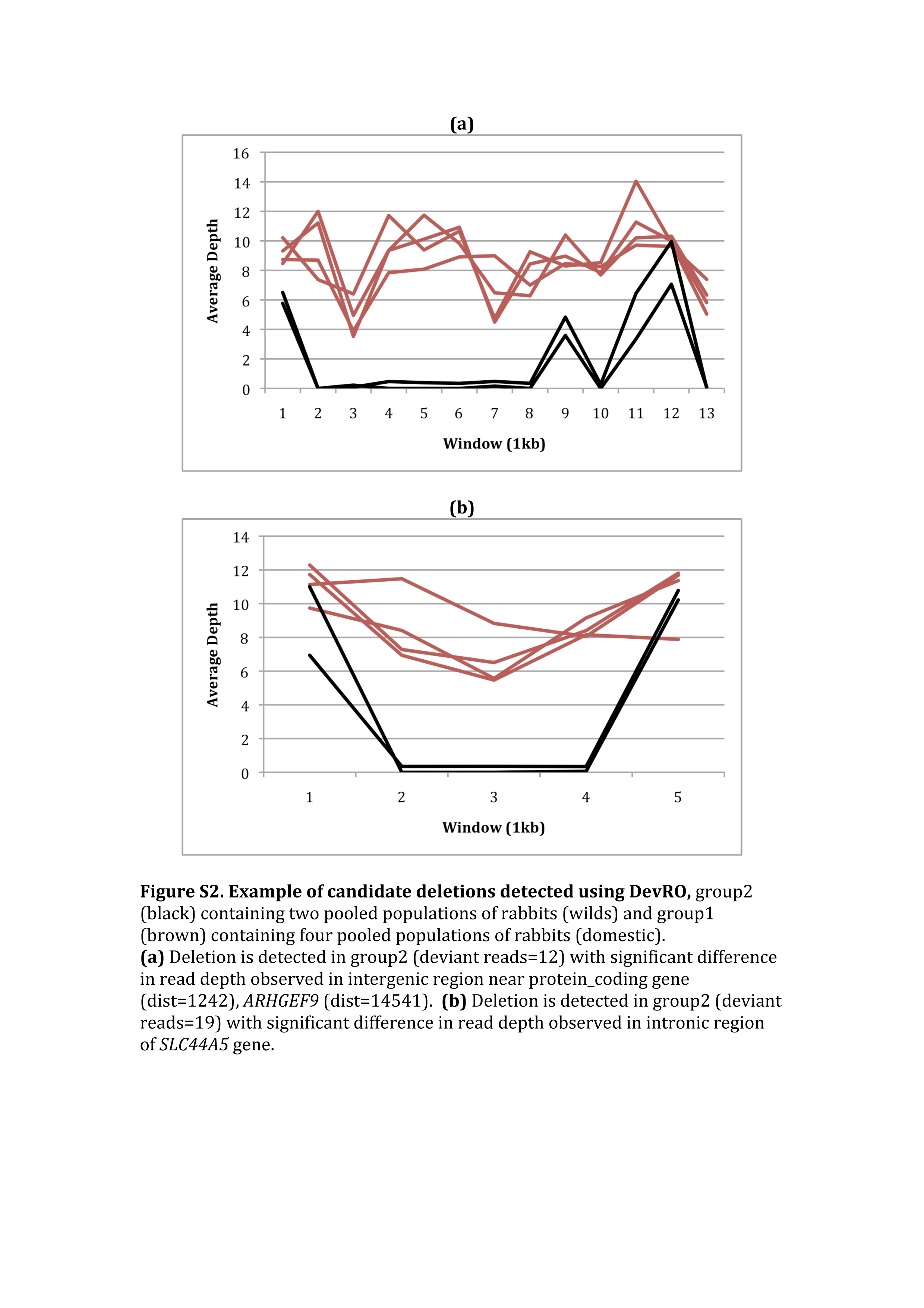
Example of candidate deletions detected using DevRO, group2 (black) containing two pooled populations of rabbits (wilds) and group1 (brown) containing four pooled populations of rabbits (domestic). **(a)** Deletion is detected in group2 (deviant reads=12) with significant difference in read depth observed in intergenic region near protein_coding gene (dist=1242), *ARHGEF9* (dist=14541). **(b)** Deletion is detected in group2 (deviant reads=19) with significant difference in read depth observed in intronic region of *SLC44A5* gene.

**Table S1.** Candidate deletions and duplications predicted by DevRO in each group.

| **Chr** | **Start** | **End** | **Size (bp)** | **Type** | **Group** |
| --- | --- | --- | --- | --- | --- |
| 1 | 10020939 | 10024805 | 3866 | Deletion | group2 |
| 1 | 28072850 | 28076964 | 4114 | Deletion | group2 |
| 1 | 35966681 | 35970344 | 3663 | Deletion | group1 |
| 1 | 40667085 | 40676380 | 9295 | Deletion | group2 |
| 1 | 47873948 | 47875998 | 2050 | Deletion | group2 |
| 1 | 47876478 | 47878705 | 2227 | Deletion | group2 |
| 1 | 54143394 | 54146991 | 3597 | Deletion | group2 |
| 1 | 71477307 | 71480636 | 3329 | Deletion | group2 |
| 1 | 101735057 | 101740562 | 5505 | Deletion | group2 |
| 1 | 101828749 | 101832131 | 3382 | Deletion | group2 |
| 1 | 109464153 | 109467763 | 3610 | Deletion | group2 |
| 1 | 110465935 | 110469799 | 3864 | Deletion | group2 |
| 1 | 111809164 | 111812815 | 3651 | Deletion | group2 |
| 1 | 114469995 | 114477656 | 7661 | Deletion | group2 |
| 1 | 118295065 | 118300976 | 5911 | Deletion | group2 |
| 1 | 126104457 | 126106943 | 2486 | Deletion | group2 |
| 1 | 127914075 | 127917518 | 3443 | Deletion | group2 |
| 1 | 134123231 | 134126802 | 3571 | Deletion | group2 |
| 1 | 141557162 | 141568087 | 10925 | Deletion | group2 |
| 1 | 143422101 | 143428148 | 6047 | Deletion | group2 |
| 1 | 147014667 | 147018084 | 3417 | Duplication | group1 |
| 1 | 147014896 | 147018485 | 3589 | Duplication | group1 |
| 1 | 149359205 | 149363430 | 4225 | Deletion | group2 |
| 1 | 152834092 | 152837418 | 3326 | Deletion | group2 |
| 1 | 155627655 | 155632178 | 4523 | Deletion | group2 |
| 1 | 162395864 | 162400029 | 4165 | Deletion | group2 |
| 1 | 164028470 | 164035995 | 7525 | Deletion | group2 |
| 1 | 167950071 | 167955986 | 5915 | Deletion | group2 |
| 1 | 179042360 | 179051729 | 9369 | Deletion | group2 |
| 1 | 188654581 | 188662331 | 7750 | Deletion | group2 |
| 2 | 8907804 | 8911624 | 3820 | Deletion | group2 |
| 2 | 29249655 | 29251880 | 2225 | Deletion | group2 |
| 2 | 36752580 | 36760767 | 8187 | Deletion | group2 |
| 2 | 52397532 | 52405411 | 7879 | Deletion | group2 |
| 2 | 63893771 | 63905895 | 12124 | Deletion | group2 |
| 2 | 77667693 | 77670961 | 3268 | Deletion | group2 |
| 2 | 83781069 | 83790995 | 9926 | Deletion | group2 |
| 2 | 135624242 | 135629669 | 5427 | Deletion | group2 |
| 2 | 142195529 | 142196096 | 567 | Duplication | group1 |
| 2 | 151892483 | 151901166 | 8683 | Deletion | group2 |
| 2 | 169841289 | 169851166 | 9877 | Deletion | group2 |
| 3 | 1716363 | 1719832 | 3469 | Deletion | group1 |
| 3 | 72563366 | 72566806 | 3440 | Deletion | group2 |
| 3 | 97268553 | 97273216 | 4663 | Deletion | group2 |
| 3 | 112684708 | 112695199 | 10491 | Deletion | group2 |
| 3 | 126363521 | 126369602 | 6081 | Deletion | group2 |
| 3 | 142492575 | 142495252 | 2677 | Duplication | group1 |
| 3 | 145155895 | 145163627 | 7732 | Deletion | group2 |
| 4 | 18742611 | 18745883 | 3272 | Deletion | group2 |
| 4 | 28982204 | 28985227 | 3023 | Deletion | group2 |
| 4 | 55327156 | 55330354 | 3198 | Deletion | group2 |
| 5 | 20859 | 24369 | 3510 | Deletion | group2 |
| 5 | 344827 | 349083 | 4256 | Duplication | group1 |
| 5 | 36303639 | 36312856 | 9217 | Deletion | group2 |
| 6 | 3363358 | 3370601 | 7243 | Deletion | group2 |
| 6 | 19228634 | 19230950 | 2316 | Deletion | group2 |
| 7 | 12981275 | 12985336 | 4061 | Deletion | group2 |
| 7 | 26803044 | 26807249 | 4205 | Deletion | group1 |
| 7 | 26803044 | 26806951 | 3907 | Deletion | group1 |
| 7 | 31718171 | 31726075 | 7904 | Deletion | group2 |
| 7 | 32507894 | 32511721 | 3827 | Deletion | group2 |
| 7 | 54843273 | 54850795 | 7522 | Deletion | group2 |
| 7 | 58958722 | 58961869 | 3147 | Duplication | group1 |
| 7 | 70261896 | 70264484 | 2588 | Duplication | group1 |
| 7 | 79489403 | 79497625 | 8222 | Deletion | group2 |
| 7 | 120402118 | 120409226 | 7108 | Deletion | group2 |
| 7 | 120749915 | 120756202 | 6287 | Deletion | group2 |
| 7 | 125173053 | 125175145 | 2092 | Duplication | group1 |
| 7 | 125386462 | 125391036 | 4574 | Deletion | group2 |
| 7 | 133758326 | 133762484 | 4158 | Deletion | group2 |
| 7 | 141972775 | 141979856 | 7081 | Deletion | group2 |
| 7 | 163088895 | 163089089 | 194 | Duplication | group1 |
| 8 | 14800377 | 14803361 | 2984 | Deletion | group1 |
| 8 | 44992852 | 44995628 | 2776 | Deletion | group2 |
| 8 | 45505862 | 45509228 | 3366 | Deletion | group2 |
| 8 | 56632623 | 56640773 | 8150 | Deletion | group2 |
| 8 | 57402830 | 57408135 | 5305 | Deletion | group2 |
| 8 | 59752725 | 59758567 | 5842 | Deletion | group2 |
| 8 | 59950628 | 59954953 | 4325 | Deletion | group2 |
| 8 | 65776341 | 65780761 | 4420 | Deletion | group2 |
| 8 | 71964830 | 71969106 | 4276 | Deletion | group2 |
| 8 | 80648555 | 80655525 | 6970 | Deletion | group2 |
| 8 | 84668351 | 84671153 | 2802 | Deletion | group2 |
| 8 | 92232111 | 92239304 | 7193 | Deletion | group2 |
| 8 | 111417475 | 111419034 | 1559 | Duplication | group2 |
| 9 | 7594534 | 7599049 | 4515 | Deletion | group1 |
| 9 | 14786168 | 14793317 | 7149 | Deletion | group2 |
| 9 | 19350787 | 19352353 | 1566 | Duplication | group1 |
| 9 | 75701994 | 75706928 | 4934 | Deletion | group2 |
| 9 | 75701994 | 75705403 | 3409 | Deletion | group2 |
| 9 | 104158641 | 104161335 | 2694 | Deletion | group2 |
| 9 | 107657364 | 107664776 | 7412 | Deletion | group2 |
| 9 | 111153173 | 111160425 | 7252 | Deletion | group2 |
| 9 | 113169200 | 113170457 | 1257 | Duplication | group1 |
| 10 | 17097046 | 17100859 | 3813 | Deletion | group1 |
| 10 | 43269463 | 43276912 | 7449 | Deletion | group2 |
| 11 | 3337013 | 3340708 | 3695 | Deletion | group2 |
| 11 | 20268944 | 20276319 | 7375 | Deletion | group2 |
| 11 | 23776245 | 23782340 | 6095 | Deletion | group2 |
| 11 | 24923632 | 24931227 | 7595 | Deletion | group2 |
| 11 | 54783931 | 54788568 | 4637 | Deletion | group2 |
| 11 | 60071092 | 60075694 | 4602 | Deletion | group2 |
| 11 | 80586769 | 80593702 | 6933 | Deletion | group2 |
| 12 | 9811249 | 9819292 | 8043 | Deletion | group2 |
| 12 | 13276045 | 13283508 | 7463 | Deletion | group2 |
| 12 | 22362392 | 22366588 | 4196 | Duplication | group2 |
| 12 | 67864698 | 67868786 | 4088 | Deletion | group2 |
| 12 | 84399800 | 84402862 | 3062 | Deletion | group2 |
| 12 | 84729252 | 84737832 | 8580 | Deletion | group2 |
| 12 | 87482467 | 87491216 | 8749 | Deletion | group2 |
| 12 | 97121922 | 97125898 | 3976 | Deletion | group2 |
| 12 | 102964911 | 102972364 | 7453 | Deletion | group2 |
| 12 | 105645122 | 105653497 | 8375 | Deletion | group2 |
| 12 | 109793243 | 109801565 | 8322 | Deletion | group2 |
| 12 | 124576676 | 124582814 | 6138 | Deletion | group2 |
| 12 | 126136891 | 126141357 | 4466 | Deletion | group2 |
| 12 | 128294878 | 128296216 | 1338 | Duplication | group1 |
| 12 | 132349837 | 132354005 | 4168 | Deletion | group2 |
| 12 | 137210541 | 137218192 | 7651 | Deletion | group2 |
| 12 | 143706074 | 143708982 | 2908 | Deletion | group2 |
| 12 | 143826021 | 143831881 | 5860 | Deletion | group2 |
| 12 | 145095491 | 145096647 | 1156 | Duplication | group1 |
| 12 | 145961895 | 145967035 | 5140 | Deletion | group2 |
| 13 | 16839514 | 16844241 | 4727 | Deletion | group2 |
| 13 | 21433995 | 21441958 | 7963 | Deletion | group2 |
| 13 | 59226582 | 59234336 | 7754 | Deletion | group1 |
| 13 | 90236204 | 90239876 | 3672 | Deletion | group2 |
| 13 | 98765308 | 98769604 | 4296 | Deletion | group2 |
| 14 | 5779535 | 5782552 | 3017 | Deletion | group2 |
| 14 | 7128562 | 7132003 | 3441 | Deletion | group2 |
| 14 | 9912909 | 9919030 | 6121 | Deletion | group2 |
| 14 | 15301386 | 15304922 | 3536 | Deletion | group2 |
| 14 | 20871985 | 20879971 | 7986 | Deletion | group2 |
| 14 | 30260260 | 30266974 | 6714 | Deletion | group2 |
| 14 | 32492338 | 32493295 | 957 | Duplication | group1 |
| 14 | 38219464 | 38227796 | 8332 | Deletion | group2 |
| 14 | 48897308 | 48904413 | 7105 | Deletion | group2 |
| 14 | 113290276 | 113301431 | 11155 | Deletion | group1 |
| 14 | 116014193 | 116018905 | 4712 | Deletion | group2 |
| 14 | 116242757 | 116245446 | 2689 | Deletion | group2 |
| 14 | 120894133 | 120896921 | 2788 | Deletion | group2 |
| 14 | 124403247 | 124406724 | 3477 | Duplication | group1 |
| 14 | 124404036 | 124407175 | 3139 | Duplication | group1 |
| 14 | 131825657 | 131829899 | 4242 | Deletion | group2 |
| 14 | 132956315 | 132960054 | 3739 | Deletion | group2 |
| 14 | 137490777 | 137495281 | 4504 | Deletion | group2 |
| 14 | 146288460 | 146294205 | 5745 | Deletion | group2 |
| 14 | 150162478 | 150170001 | 7523 | Deletion | group2 |
| 14 | 151553461 | 151555787 | 2326 | Deletion | group2 |
| 14 | 151724087 | 151732320 | 8233 | Deletion | group2 |
| 14 | 153976963 | 153982969 | 6006 | Deletion | group2 |
| 14 | 154545468 | 154549225 | 3757 | Deletion | group2 |
| 15 | 26808869 | 26811837 | 2968 | Deletion | group2 |
| 15 | 30971660 | 30978614 | 6954 | Deletion | group2 |
| 15 | 30972230 | 30978614 | 6384 | Deletion | group2 |
| 15 | 40544197 | 40550341 | 6144 | Deletion | group2 |
| 15 | 61930692 | 61940459 | 9767 | Deletion | group2 |
| 15 | 63575127 | 63579020 | 3893 | Deletion | group2 |
| 15 | 63643499 | 63650867 | 7368 | Deletion | group2 |
| 15 | 65003580 | 65006277 | 2697 | Duplication | group1 |
| 15 | 72690503 | 72693247 | 2744 | Deletion | group2 |
| 15 | 78477184 | 78481021 | 3837 | Deletion | group2 |
| 15 | 80232545 | 80267536 | 34991 | Duplication | group2 |
| 15 | 80233369 | 80267734 | 34365 | Duplication | group2 |
| 15 | 82895900 | 82901437 | 5537 | Deletion | group2 |
| 15 | 86189640 | 86197057 | 7417 | Deletion | group2 |
| 15 | 96732790 | 96733582 | 792 | Duplication | group1 |
| 15 | 98159531 | 98162919 | 3388 | Deletion | group2 |
| 15 | 103898002 | 103902748 | 4746 | Deletion | group2 |
| 16 | 18033252 | 18100872 | 67620 | Deletion | group1 |
| 16 | 19590590 | 19592339 | 1749 | Duplication | group1 |
| 16 | 29964075 | 30118260 | 154185 | Duplication | group2 |
| 16 | 51032152 | 51035559 | 3407 | Deletion | group2 |
| 16 | 55597969 | 55605702 | 7733 | Deletion | group1 |
| 17 | 40281181 | 40283868 | 2687 | Deletion | group2 |
| 17 | 46968775 | 46975089 | 6314 | Deletion | group2 |
| 17 | 59950714 | 59959230 | 8516 | Deletion | group2 |
| 17 | 75224247 | 75229589 | 5342 | Deletion | group2 |
| 17 | 81264203 | 81266865 | 2662 | Deletion | group1 |
| 18 | 4310271 | 4317974 | 7703 | Deletion | group2 |
| 18 | 56156738 | 56160292 | 3554 | Deletion | group2 |
| 18 | 58650087 | 58658467 | 8380 | Deletion | group2 |
| 19 | 34305013 | 34312929 | 7916 | Deletion | group2 |
| X | 722954 | 726552 | 3598 | Deletion | group2 |
| X | 1773782 | 1781304 | 7522 | Deletion | group2 |
| X | 20611770 | 20618218 | 6448 | Deletion | group2 |
| X | 41695980 | 41698714 | 2734 | Duplication | group1 |
| X | 42248861 | 42261352 | 12491 | Deletion | group2 |
| X | 44633371 | 44637590 | 4219 | Deletion | group2 |
| X | 74054891 | 74060468 | 5577 | Deletion | group2 |
| X | 76516186 | 76524291 | 8105 | Deletion | group2 |
| X | 86022441 | 86030686 | 8245 | Deletion | group2 |
| X | 96399403 | 96403985 | 4582 | Deletion | group2 |
| X | 96673270 | 96679147 | 5877 | Deletion | group2 |
| X | 108480223 | 108486844 | 6621 | Deletion | group1 |

**Table S2.** Subset of candidate Inversions detected by DevRO not in SVDetect list.

| **Chromosome** | **Start** | **End** | **Group<sup>a</sup>** | **Size (bp)** |
| --- | --- | --- | --- | --- |
| 9 | 18388001 | 18388750 | group2 | 749 |
| 8 | 729001 | 729787 | group2 | 786 |
| 13 | 13931001 | 13931791 | group1 | 790 |
| 16 | 13903001 | 13903829 | group2 | 828 |
| 14 | 30506001 | 30506889 | group1 | 888 |
| 9 | 37406000 | 37406906 | group1 | 906 |
| 4 | 45058000 | 45058929 | group1 | 929 |
| 4 | 37565000 | 37565961 | group2 | 961 |
| 13 | 80351000 | 80351992 | group2 | 992 |
| 13 | 120546000 | 120547010 | group1 | 1010 |
| 14 | 98774000 | 98775037 | group1 | 1037 |
| 14 | 86379000 | 86380041 | group1 | 1041 |
| 14 | 123838000 | 123839049 | group1 | 1049 |
| 16 | 29365001 | 29366168 | group1 | 1167 |
| 3 | 22743001 | 22744190 | group1 | 1189 |
| 4 | 37575000 | 37576609 | group1 | 1609 |
| 14 | 30518001 | 30519796 | group1 | 1795 |
| 16 | 27289001 | 27290858 | group2 | 1857 |
| 11 | 8202001 | 8203947 | group1 | 1946 |
| 3 | 22682001 | 22684150 | group2 | 2149 |
| 4 | 35718000 | 35720375 | group2 | 2375 |
| 14 | 30442001 | 30444616 | group1 | 2615 |
| 2 | 172979000 | 172981685 | group1 | 2685 |
| 9 | 48563000 | 48565730 | group1 | 2730 |
| 16 | 29574001 | 29576732 | group2 | 2731 |
| 3 | 21551001 | 21553879 | group1 | 2878 |
| 14 | 98958000 | 98961458 | group2 | 3458 |
| 4 | 40112000 | 40115630 | group1 | 3630 |
| 3 | 22216001 | 22219761 | group1 | 3760 |
| 16 | 52641000 | 52644941 | group2 | 3941 |
| 13 | 13506001 | 13510568 | group2 | 4567 |
| 4 | 40116000 | 40120949 | group1 | 4949 |
| 8 | 43127000 | 43131968 | group1 | 4968 |
| 9 | 15468001 | 16079438 | group1 | 611437 |
| 16 | 19914001 | 20607842 | group1 | 693841 |
| 11 | 37666000 | 46273752 | group2 | 8607752 |

<sup>a</sup> Group based on test data representing domestic or wild

**Table S3.** Simulation Analysis for deletion in the reference assembly

| locusID | coordinates<sup>a</sup> | size(bp) | region info<sup>b</sup> | 5' mapability<sup>c</sup> | 3' mapability<sup>d</sup> | DevRO BP <sup>e</sup> | ReadCounts<sup>f</sup> |
| --- | --- | --- | --- | --- | --- | --- | --- |
| region1 | chr1:10,974,545-11,004,544 | 30000 | lots of repeats | 0.254498332 | 0.991055897 | 10974446 | 32 |
| region2 | chr1:20,976,760-20,977,759 | 1000 | simple tandem repeat | 0.876914441 | 0.992240837 | 20976370 | 160 |
| region3 | chr1:50,989,028-50,991,029 | 2000 | LINES;SINES;simple | 0.982284416 | 0.937495072 | 50989291 | 39 |
| region4 | chr1:40,252,552-40,257,556 | 5005 | No repeats, only masked ones | 0.982258048 | 0.987218825 | 40254644 | 24 |
| region5 | chr1:121,020,709-121,027,708 | 7000 | No Repeats | 0.996043815 | 0.995074997 | 121020479 | 22 |
| region6 | chr1:131,052,703-131,062,702 | 10000 | repeats, LIR, simple | 0.994097772 | 0.93532862 | 131055431 | 183 |
| region7 | chr1:8,062,089-8,067,088 | 5000 | No overlapping repeat | 0.960199177 | 0.978494121 | 8066956 | 23 |
| region8 | chr1:28,400,306-28,405,305 | 5000 | LINE on one side of breakpoint | 0.980429893 | 0.000427342 | 28400443 | 22 |
| region9 | chr1:58,030,839-58,100,838 | 70000 | lots of repeats | 0.007576796 | 0.992102998 | NA | NA |
| region10 | chr1:3,029,339-3,032,338 | 3000 | SINE, with no repeats at bp | 0.992139649 | 0.973058135 | 3029709 | 46 |

<sup>a</sup> coordinates of simulated deletions<sup>b</sup> Information of overlapping repeats at breakpoints (Repeat information extracted from UCSC Santa Cruze Browser)<sup>c</sup> 5'-1000 weighted mean mapability<sup>d</sup> 3'+1000 weighted mean mapability<sup>e</sup> Breakpoint predicted by DevRO for simulated deletions<sup>f</sup> Number of read counts supporting the simulated deletion

